# Effect of Poli I:C in murine model of autoimmune thyroiditis

**DOI:** 10.1101/034066

**Authors:** C Vera, M. Barria

## Abstract

Hashimoto’s thyroiditis is one of the most common autoimmune diseases in humans and, similar to other autoimmune diseases, is multifactorial in nature. Moreover, the expression of TLRs has been implicated in the pathogenesis of autoimmune and inflammatory diseases, such as diabetes and insulinitis. The TLRs are a family of at least 10 receptors associated with innate immunity that are present in monocytes, macrophages, and dendritic cells, which recognize highly conserved patterns of the surface of microorganisms, such as LPS, peptidoglycan, and dsRNA and CpG sequences.

The aim of this study was to evaluate mRNA of the TLR3, TLR9 and cytokines (IFN-beta, TNF and IL-1) in thyroid follicular cells obtained of mice with autoimmune experimental thyroiditis (EAT). For that C57BL/6 mice were immunized with Tg plus complete Freund’s adyuvant and treated or not with Poli I:C. Anti-thyroglobulin autoantibodies level was determined by indirect ELISA. Also isolated thyroid follicles were treated *in vitro* with Poli IC. The results show that TLR3 receptors and some cytokines (IFN-beta, TNF and IL-10) are more highly expressed in thyroid follicular cells treated with Poli I:C. and in the animals with TAE, which present humoral autoimmune response. Taken together, these results suggest that TLR3 expression in thyroid follicular cells induces signaling mechanisms to enhanced IL-1, TNF and IFN-β levels during pathogen infections and that could trigger of autoimmune thyroiditis.

## INTRODUCTION

Hashimoto’s thyroiditis is one of the most common autoimmune diseases in humans and the most common cause of hypothyroidism in areas of the world where iodine concentrations are sufficient, in some countries may reach 5% of the population. It is an organ-specific autoimmune disorder characterized by cellular and humoral autoimmunity that causes a progressive failure of the thyroid gland due to extensive infiltration of T and B lymphocytes, plasma cells and macrophages, production of a high amount of anti-peroxidase, anti-thyroglobulin and anti-TSH-R auto-antibodies, which generate progressive destruction of the organ (Carmody and Chen, 2007). It is a disorder more prevalent in women than in men (ratio of 10: 1 to 20: 1) and between 45 to 65 years. The development of the disease requires the coordinated expression of multiple immuno-related genes capable of encoding various molecules. There is an increased expression of MHC-I and MHC-II molecules in thyrocytes of patients, which suggests they may have a role as antigen presenting cells (Giuliani et al., 2010). It has also been observed expression of adhesion molecules ICAM-1, B7–1, costimulatory molecules essential for interaction of immune cell, INF, IP-10 protein (protein which is induced by IFN-γ and ligand for the CXC chemokine with chemotactic activity on lymphoid cells) and FAS genes, which encode a receptor containing a death domain that interact with their ligands on the surface of thyrocytes could induce apoptosis and be a possible cause of destruction of the gland (García-López et al., 2001; Pesce et al., 2002; Giordano et al., 1997; Pearce et al, 2003). It has further demonstrated that human thyroid follicular cells chemotactic factor produced IL-16, capable of participating in lymphocyte recruitment during inflammatory process (Gianoukaki AG. Et al., 2003). Moreover, these cells can be target of LPS, which induce the expression of several chemokines (MIP-3, MCP-1, TARC) facilitating lymphocyte infiltration (Damotte D. et al., 2003).

Despite the findings, it has not yet been demonstrated that these cells alone can trigger an autoimmune response (Margolese, H et al., 1994; Maile R et al., 2000; YS Li. et al., 2004). It is known that thyrocytes play a central role in the thyroid pathogenic process, but it is not clear whether gene expression is an a key activity of thyrocytes to induce disease or a secondary response to cytokines produced by infiltrating immune cells. The molecular events involved are unknown as well; it is known that most of the genes expressed are regulated by transcription factors, where NF-κB activated via TLR has an important role in activating thyroid follicular cells (Barría et al., 2005, manuscript). It described the production of proinflammatory cytokines induced by this mechanism (IL-1, TNF-α and IL-6) are able to modulate thyroid hormonogenesis *in vitro* after receptor activation by PAMPs (Yamazaki K et al., 2007).

Human thyrocytes surrounded by immune cells in Hashimoto’s thyroiditis show TLR3 sobreexpesión (Harii, N et al., 2005). It has also found TLR9 expression in thyrocytes, but little is known about their functionality.

While the cause of autoimmune damage generated in autoimmune thyroiditis is unknown, the viral infection would have an important role as disease trigger, considering that the dsRNA produced during viral replication of many viruses is a potent inducer of IFN-1 and dendritic cell activator (Takeda et al, 2003). An example of infectious agents involved in the induction of thyroiditis autoimmune is the association between infection and Foamy Virus Quervain’s thyroiditis. Harry N. et al. in 2005, demonstrated functional TLR3 overexpression in a cell line of rat thyroid (FRTL-5) after stimulating the receptor with Poly I: C, inducing the expression of pro-inflammatory genes via activation of NF-κB. They also observed the overexpression TLR3 in human thyrocytes cultured transfected dsRNA and in thyroid gland biopsies from patients with Hashimoto’s thyroiditis.

Toll-like receptors (TLRs) are a family of receptors that recognize pathogen-associated molecular patterns (PAMPs) as ligands, which include molecules of gram positive and gram negative bacteria, viral nucleic acids, fungi, protozoa (Takeda et al., 2007; Kawai et al., 2007) as well as a variety of host-derived (Li et al., 2009). It has been detected between ten and fifteen TLR receptors in mammals, being identified eleven in human and thirteen in mice. For example, TLR3 recognizes double-stranded RNA (dsRNA) viral and its action can be simulated using the synthetic analogue referred Polyinosinic : polycytidylic acid or Poly (I:C) (Alexopoulou et al, 2001); while TLR9 is stimulated by unmethylated CpG motifs which is present in bacterial and viral DNA, whose effect can be simulated with synthetic oligodeoxynucleotides with CpG motifs identical (Hemmi et al reasons., 2000; Klinman et al., 1996; Krieg et al. 1995). TLRs are expressed primarily in immune cells, namely in macrophages, dendritic cells, neutrophils, NK cells, B and T lymphocytes; but they are also present in other cell types such as endothelial cells, fibroblasts and epithelial cells of different organs (prostate, ovary, trachea, bronchus, lung, placenta, liver, spleen, thyroid, etc.), either constitutively or induced by infection.

The innate immune system is activated upon recognition of PAMPs, triggering the release of proinflammatory cytokines by activation of NF-κB and other signaling pathways (Kumar et al, 2009; Foster SL et al, 2009) and participating in the development of adaptive immune responses by stimulating costimulatory molecules of antigen presenting cells (Takeda K et al., 2003). It is conjectured that individual particular responses to this activation mediated by TLRs could break tolerance to self and play a crucial role in the susceptibility to develop autoimmune diseases such as rheumatoid arthritis, type 1 diabetes mellitus, lupus erythematosus, Hashimoto’s thyroiditis, among others (Leadbetter EA et al., 2002, Waldner, 2009; Patole PS et al., 2005; Zipris D et al., 2005; Li M et al., 2009; Xiangrong y col, 2011).

The main theory to explain how autoantigens are generated and how they are able to break tolerance is supported both antigenic similarity between exogenous and endogenous ligands recognized by the same TLR, as well as the idea that autoantigens are such because are autoadyuvantes, that is, can activate the innate immune system directly promoting autoimmune responses. It is known that the same receptors that distinguish microbial TLR ligands with strong adjuvant activity, also they recognize molecules released in response to cellular damage, apoptotic clearance of debris and tissue repair; For example, TLR3 recognizes viral dsRNA and is also a sensor endogenous tissue necrosis. This antigenic similarity under certain circumstances could induce the activation of dendritic cells, macrophages and other antigen presenting cells, triggering autoreactive response mediated by B and T lymphocyte. This is reflected in the high proportion of observable autoantibodies in autoimmune diseases (lupus erythematosus, scleroderma, Sjögren’s syndrome, uveitis, etc.), which react with the DNA, RNA or macromolecular complexes containing DNA or RNA. (Fang et al, 2010;. Marshak-Rothstein, 2006).

In the 90s it has been suggested that the autoinmune activation by a viral agent can cause local tissue infection, abnormal induction or enhanced expression of MHC genes with the consequent autoantigen presentation to immune cells and T cell activation (Benoist et al., 1998; Horwitz et al., 1998; Wekerle et al., 1998). Current results suggest that various microbial infections may contribute to the pathogenesis of diseases such as uveitis. It was observed that signaling via TLR2, TLR3, TLR4 and TLR9 was highly redundant in the adjuvant effect needed to induce experimental autoimmune uveitis when each agonist was able to increase the levels of disease in a regime of immunization using complete adjuvant Freund (Fang et al, 2010).

*In vitro* and *in vivo* studies show important link between the immune complexes containing DNA and RNA, activation of TLRs with subsequent expression of type I interferon and proinflammatory cytokines and induction of an adaptive immune response. *In vitro* studies showed that the TLR3 agonist, Poly I:C, induced synthesis of IL-17A and IL-21 in human CD4+ T cells, which points to an important role of TLR3 in the regulation of CD4 + T cells in autoimmunity (CK Holm et al., 2009). It has been seen that Poly I: C increases the severity of experimental autoimmune uveitis in association with increased levels of Th1/Th17 in mice, delayed-type hypersensitivity responses and proliferation of Ag-specific T cells. In addition, the production of IL-17 and IFN-γ by Ag-specific lymph node cells was markedly increased in the group treated with Poly I: C (Xiangrong R et al, 2011.). With regard to TLR9, studies in animals deficient in this receptor have been controversial and it is unclear whether it plays a key role in the development of autoimmune responses. *In vitro* data involves FcγRs-mediated release of immune complexes formed with nucleic acids into dendritic cell compartments containing TLR9 as an important event in the pathogenesis of systemic lupus erythematosus. These immune complexes also activate monocytes and neutrophils contribute to the inflammatory process and/or regulatory pathways operating in autoimmune diseases. (Marshak-Rothstein, 2006).

Previous studies in our laboratory showed detectable amounts of TLRs (mRNA and protein) in thyrocytes from control animals and increased expression over time in mice with experimental autoimmune thyroiditis. This can be attributed to an activation and transformation of thyrocytes in antigen presenting cells through the inflammatory process or mediated by autoantibodies, phenomenon that would be involved in the development and progression of autoimmune thyroiditis (Swain et al, 2005).

This study plans to analyze the ability of microbial analogues Poly I: C, bacterial DNA and DNA with unmethylated CpG motifs to increase the TLR3, TLR9, INF-β, TNF and IL-1 mRNA expression level in follicular thyroid cells from mice with EA, who may be involved in the development and progression of the disease. Furthermore, it evaluates the possible adjuvant effect of Poly: C in the induction of experimental autoimmune thyroiditis.

## MATERIALS AND METHODS

### Animals

Female mice Rockefeller strain of 2–3 months of age, maintained with food and water freely available, belonging to the Institute of Immunology, Universidad Austral de Chile, were used in all experiments.

### Antigen and reagents

Thyroglobulin murine was used to induce experimental autoimmune thyroiditis., which were obtained from normal mouse thyroid glands by mechanic homogenization and sonication (Ultrasonic Homogenizer 4710 Series, Cole Parmer) followed by salt precipitatation with 45% ammonium sulfate (Merck). Finally protein pellet was resuspended in phosphate saline buffer pH 8.0. Dialysis was carried out in saline at 4°C with three changes every 24 hours and then the solution was filtered using 0.22 micron GP Millex filters (Millipore). The concentration of total protein was quantified by UV spectrophotometry at 280 nm. (UV-120-02 Spectrophotomer, Shimadzu) and thyroglobulin purity was checked by polyacrylamide gel electrophoresis in denaturing conditions. The solution was stored at 4°C.

The Freund complete adjuvant (FCA) and Freund incomplete adjuvant (FIA) were from Sigma, those used with thyroglobulin solution in a 1: 1 v/v.

Poly I:C stock solution (Sigma) was prepared with sterile PBS at a concentration of 5 mg/ml. For *in vivo* treatment was used at 50 ug/100 ul PBS, whereas for *in vitro* treatment of thyroid follicular cells in culture, was used at 50, 100 and 250 ug/ml.

### Induction of experimental autoimmune thyroiditis (EAT) and scoring

Rockefeller mice were immunized with 80 ug of Tg emulsified in FCA, subcutaneously injecting a total volume of 100 ul emulsion/mouse. The procedure was repeated a week later with FIA. As negative control, animals were injected with PBS.

EAT induction was evaluated determining the anti-Tg antibodies levels by indirect ELISA using sera obtained from immunized and unimmunized mice. In 96 well flat bottom microplates (Pierce Chemical Co.) was adhered the antigen after incorporating thyroglobulin 10 ug/ml in carbonate-bicarbonate buffer 0.05 M pH 9.6 adding 100/well and incubate overnight at 4°C. For blocking was used skim milk 5% in PBS adding 200 ul/well and incubating at 37°C for 2 hours, followed by three washes with PBS-Tween 20 (Sigma) 0.05%. Then, 100 ul/well of goat anti-mouse IgG conjugated to peroxidase antibody (Sigma) was added diluted 1:2000 in PBS/0,5% milk; was incubated one hour at 37 ° C and washed three times with PBS - Tween 20 0,05%.

Finally, the substarte used was OPD ortho-phenylenediamine (Sigma)(0.4 mg/ml) at phosphate citrate buffer 0.15 M pH 5.0 in presence of hydrogen peroxide 0,5% v/v. The incubation was at 37°C for 10 min. and the colorimetric reaction was stopped with 50 u/well of H2SO4 2M. The autoantibodies titer is determined measuring spectrophotometrically absorbances at 490 nm with an ELISA reader (Labsystems Uniskan I) against a blank containing substrate more H2SO4 2M. For animals immunized was considered values absorbance between 0.5 – 1.0. Each analysis was done in duplicate.

### Primary culture of thyroid follicles

Thyroid follicles cultures from mice thyroid glands were used in the study. Thyroids were removed aseptically and placed in sterile Petri dish with incomplete RPMI 1640 medium (Gibco), supplemented with Amphotericin B (HyClone) 1% and Penicillin-Streptomycin (Biological Industries) 1%. In laminar flow chamber and under the microscope, the loose connective tissue was extracted and then these glands were dissected with 21G needle in 10 to 12 fragments. For enzymatic digestion, they were immersed in 1 ml digestion buffer containing 112 U / ml collagenase type II (US Biological) and 1.2 U / ml dispase (Stemcell) dissolved in culture medium Nu-Serum (Nu-Serum IV^®^ Becton Dickinson), with continuous stirring at 3 g for 1 hour at 37°C, followed by centrifugation at 160 g. The supernatant was removed and the thyroid follicles contained in the pellet was resuspended in Nu-Serum^®^ for its incorporation into 24 wells culture dishes. Incubation was at 37 ° C, 5% CO2.

### Treatment *in vitro* and in vivo with Poly I: C

The anti-Tg antibodies levels were determinate by ELISA allowed define the two groups considered experimental studies: animals immunized and non-immunized animals.

For *in vitro* analysis, thyroid follicles suspension pooled from three immunized mice and another suspension from a pool of three non-immunized mice, each in 300 ul of complete medium Nu-Serum was prepared. Both cultivation was initially for 24 hrs, after which the original medium was removed and replaced with the same medium added with Poly I:C (50, 100 and 250 ug/ml) and incubated for further 8 hours. As controls we used only Nu-Serum medium.

In the *in vivo* study, 8 groups of 6 animals each were considered. Half of the groups were injected intraperitoneally with 50 ug of Poly I: C/100 ul of PBS to each mouse for 16 days, while the other half were injected with PBS only (control group). The second day started this process began with the immunization program to induce EAT in animals inoculated with Poly I: C and animal control; Tg plus adjuvant is used only in the first group each condition, while the other three groups are controls and adjuvant is administered, Tg and PBS, respectively. At 17° day the thyroid glands of 3 mice per group are extracted to perform thyroid follicles cultures. At day 24 the process is repeated with the remaining 3 mice.

### RNA isolation and gene expression analysis

Total RNA thyroid follicles in culture were extracted using the Σ.ZNA^®^ kit (Omega Biotek). Quantification and quality of the RNA was determined using a spectrophotometer Nano Quant (Tecan), its integrity was checked by agarose gel 1.5% and 1 ug was subjected to reverse transcription to obtain cDNA using Impront II^®^ kit (Promega).

Conventional PCR was performed using Go Taq 2X Green Master Mix kit (Promega) with specific primers for TLR3, TLR9, INF-β and IL-1 genes. Them expression was normalized to the housekeeping gene GAPDH. The relative semiquantification was performed by densitometric analysis of the PCR products after electrophoretic run on agarose gels, using the ImageJ software. The numerical values allowed to calculate the ratio between the intensity of the genes of interest and GAPDH to normalize the expression. Based on Primer-BLAST (http://www.ncbi.nlm.nih.gov/nuccore/?term=) were design the primers. The following primer sequences used are as follows:

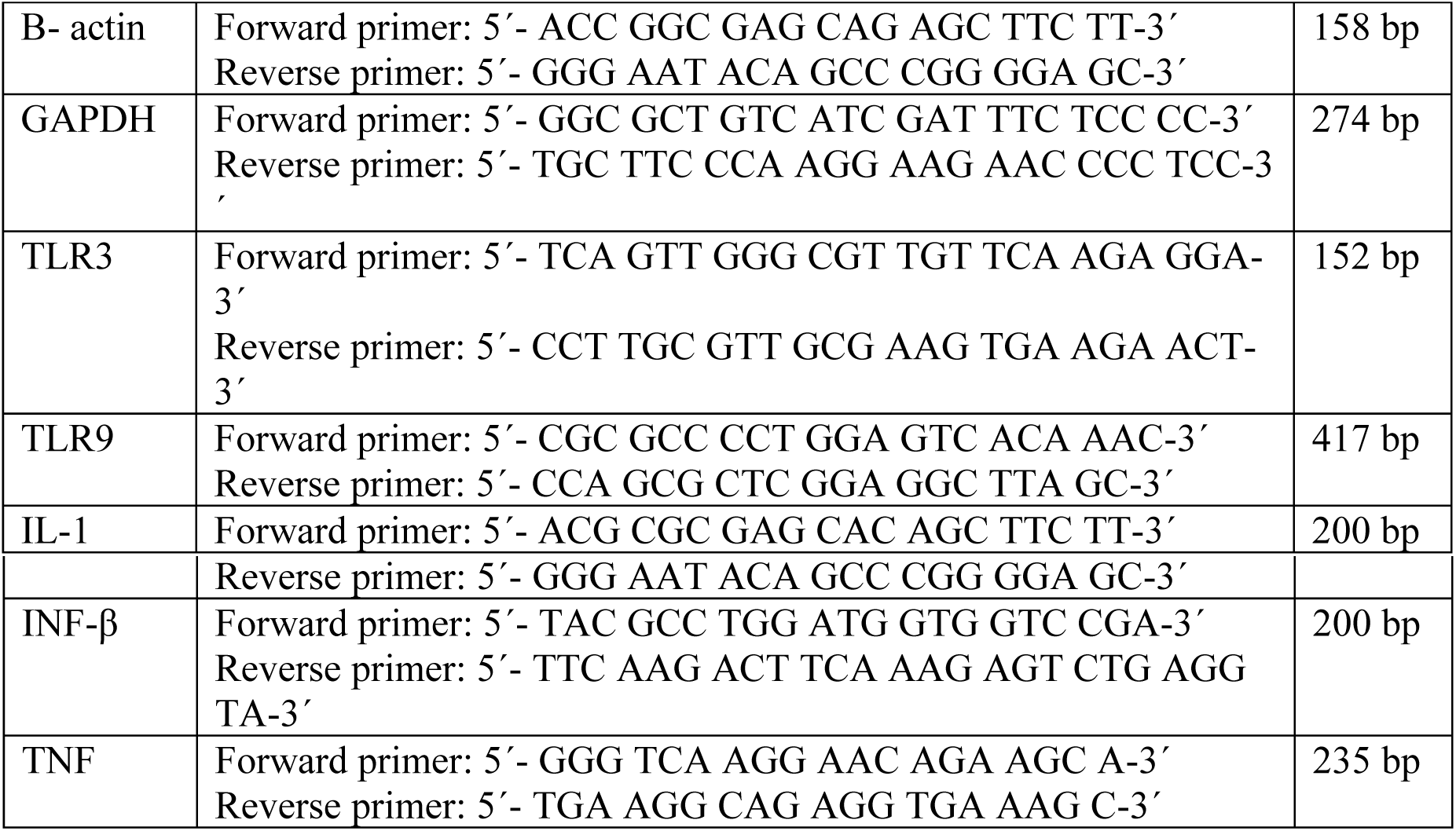

### Statistical analysis

For statistical analysis, the data obtained in this study were performed in triplicate and expressed as average with corresponding standard deviation. Simple ANOVA was used for comparison between groups and established the significance value was P <0.05. The results were plotted using SigmaPlot program 11.

## RESULTS

### Autoantibody levels in animals with experimental autoimmune thyroiditis (EAT) induced

EAT induction in experimental animals immunized with thyroglobulin (Tg) in Freund’s adjuvant was verified by determining the antibodies anti-Tg levels through an indirect ELISA, considered for this absorbance values between 0.5 - 1.0 (immunized animal). Control animals were injected with PBS. Each analysis was performed in duplicate (Figure 1). It was observed an increase in the antibodies levels in immunized animals compared to non-immunized ones at day twenty-one and twenty-eight post-immunization, showing up a slight decrease at 28 days.

**Figure 1:**
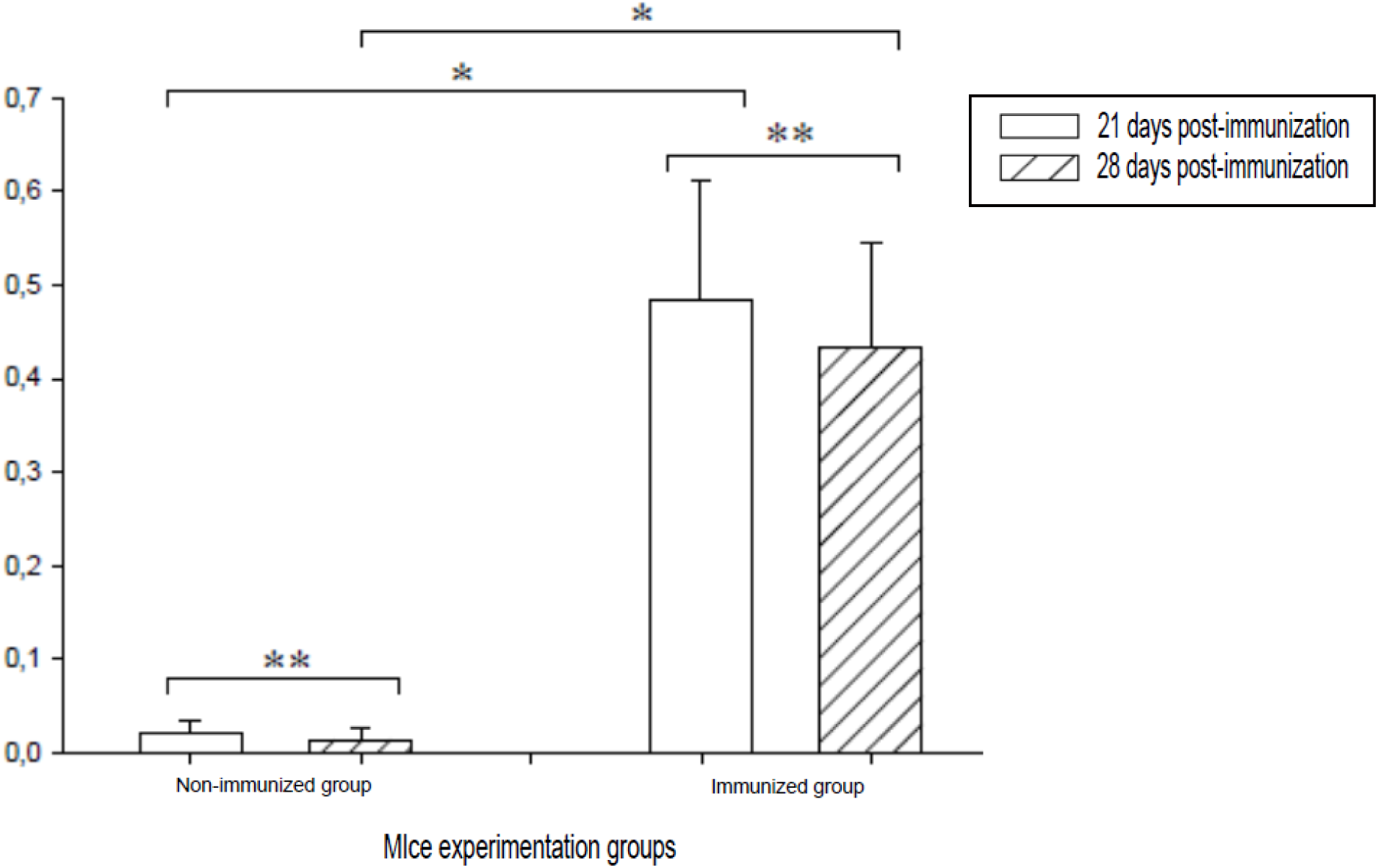
Antibodies level anti-Tg expressed as absorbance +/− S.E. in Rockefeller females mice groups non-immunized and immunized through indirect ELISA method, diluted 1:100 after the first bleeding at 21 days post-immunization and the second bleed at 28 days after the first immunization. p<0,05 (*), p>0,05 (**).

### Primary culture isolated thyroid follicles

Thyroid follicles obtained from experimental animals were cultured isolately in culture dishes. In Figure 2, thyroid follicles to 10x and 40x magnification, untreated and stimulated in vitro with Poly I:C is showed. Clearly only well preserved follicular structures on the plate is observed after digestion and disintegration of follicles (t0) and after 18 hours of incubation (t18), they show extensions and have a strong attachment to the matrix. No morphological differences were observed between the untreated follicles and stimulated with viral analog after 8 hours of incubation.

**Figure 2:**
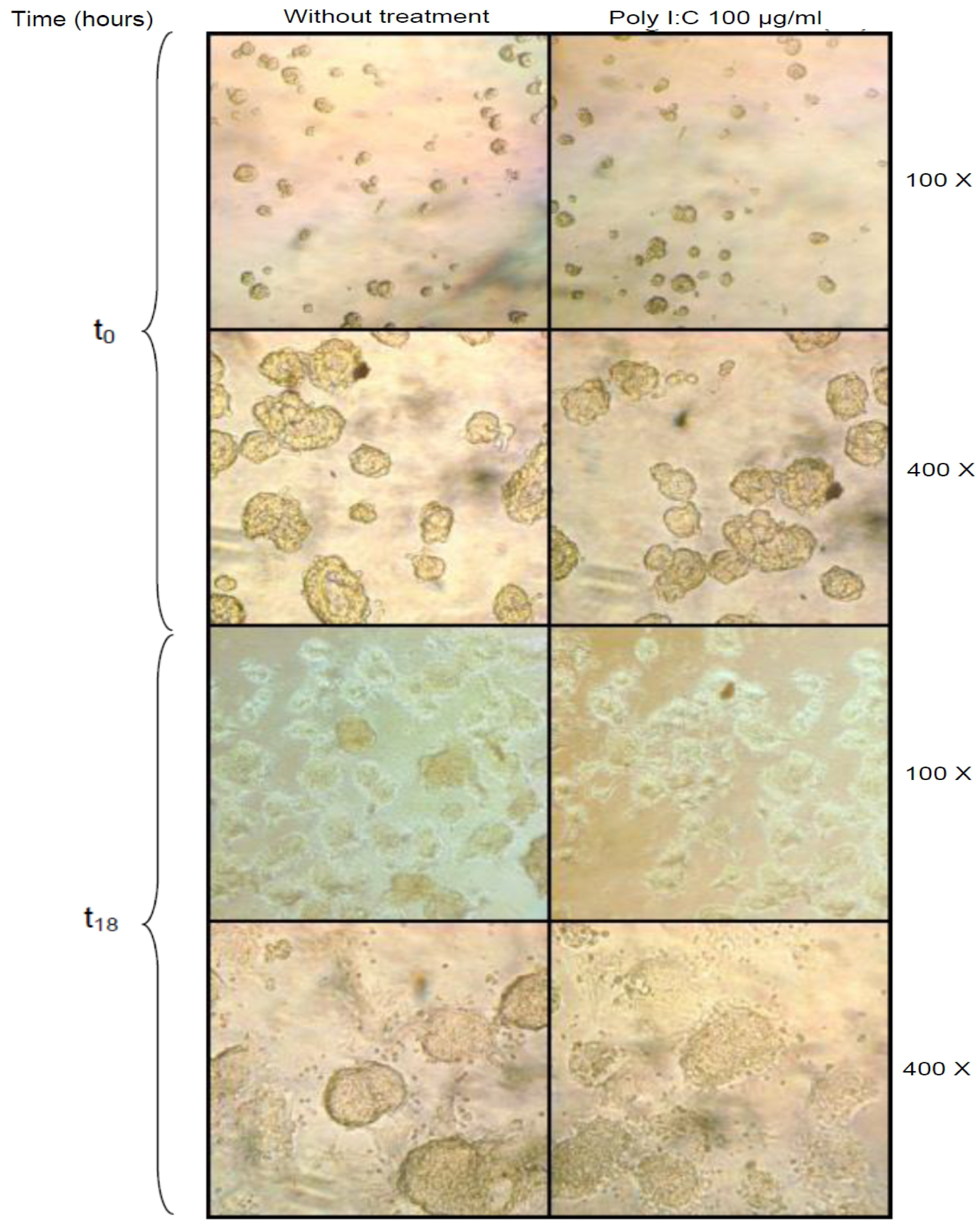
Primary culture of thyroid follicles of mice RK females; untreated and stimulated in vitro with 100 ug / ml poly (IC), at different times and optical magnification (100x and 400x).

### TLR3, TLR9, IFN-beta, IL-1 and TNF mRNA expression levels study in thyroid follicular cells from non-immunized and immunized mice treated *in vitro* with Poly I: C by semiquantitative RT-PCR

To evaluate the effect on TLR3, TLR9, INF-beta, IL-1 and TNF mRNA expression against the Poly I:C stimulation in culture, thyroid follicular cells from non-immunized and immunized animals were incubated with the viral synthetic analogous at concentrations of 0 (control), 50, 100 and 250 ug/ ml for 8 hours. These experiments were performed with other incubation times, with no significant differences in mRNA expression.

Figures 3A and 3B show TLR3 mRNA amplicons in thyrocytes from non-immunized and immunized mice, respectively, while figures 3C and 3D corresponding to GAPDH. In Figure 3E, the TLR3 expression levels under both conditions is plotted after amplicons densitometric analysis normalized with GAPDH. It can be seen that eather in immunized and non-immunized animals have a growing increase in the expression level relative to the control (no incubation with Poly I: C) when stimulated with 50 and 100 ug/ml of the analog, producing in both cases significant statistically increases. However, in non-immunized mice can be seen that stimulation with 250 ug/ml produces an abrupt decrease in mRNA expression when is compared with other concentrations and even becomes less than the control, this inhibitory effect does not occur in immunized animals when it is observed at a similar expression level to that induced by the concentration of 100 μg/ml.

**Figure 3:**
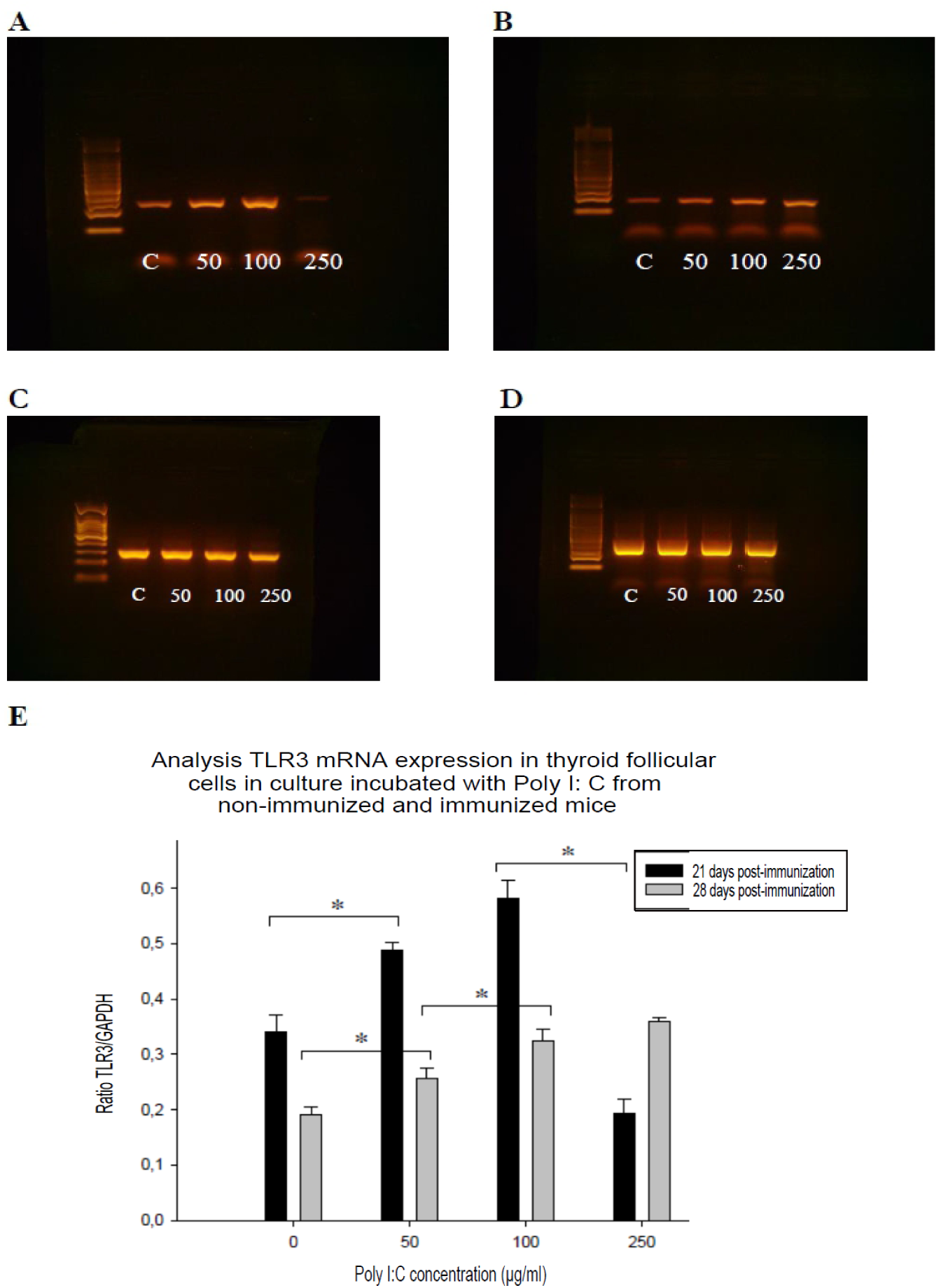
Agarose gels showing PCR-amplified fragments from TLR3 mRNA (152 bp) (**A** and **B**) and GAPDH (274 bp) (**C** and **D**) in thyroid follicles from a pool of 3 mice developed for each of the conditions: non-immunized (**A** and **C**) and immunized (**B** and **D**); which were incubated in cell culture with Poly I: C for 8 hours. **E:** Plot showing TLR3 mRNA expression ratio GAPDH-normalized. C = Control, 50 = Poli I:C 50 ug / ml, 100 = Poli I:C 100 mg/ml, 250 = Poli I:C 250 mg/ml. n = 3. p <0.05 (*), p> 0.05 (**).

In Figure 4 it is shown TLR9 and GAPDH mRNA amplification fragments in thyroid follicles from non-immunized and immunized mice stimulated in culture with Poly I:C at same concentrations and conditions given above, where the synthetic analogue did not generate major changes in the mRNA expression compared with control in either studied conditions; it can only been seen a slight reduction in the concentration 250 mg/ml.

**Figure 4:**
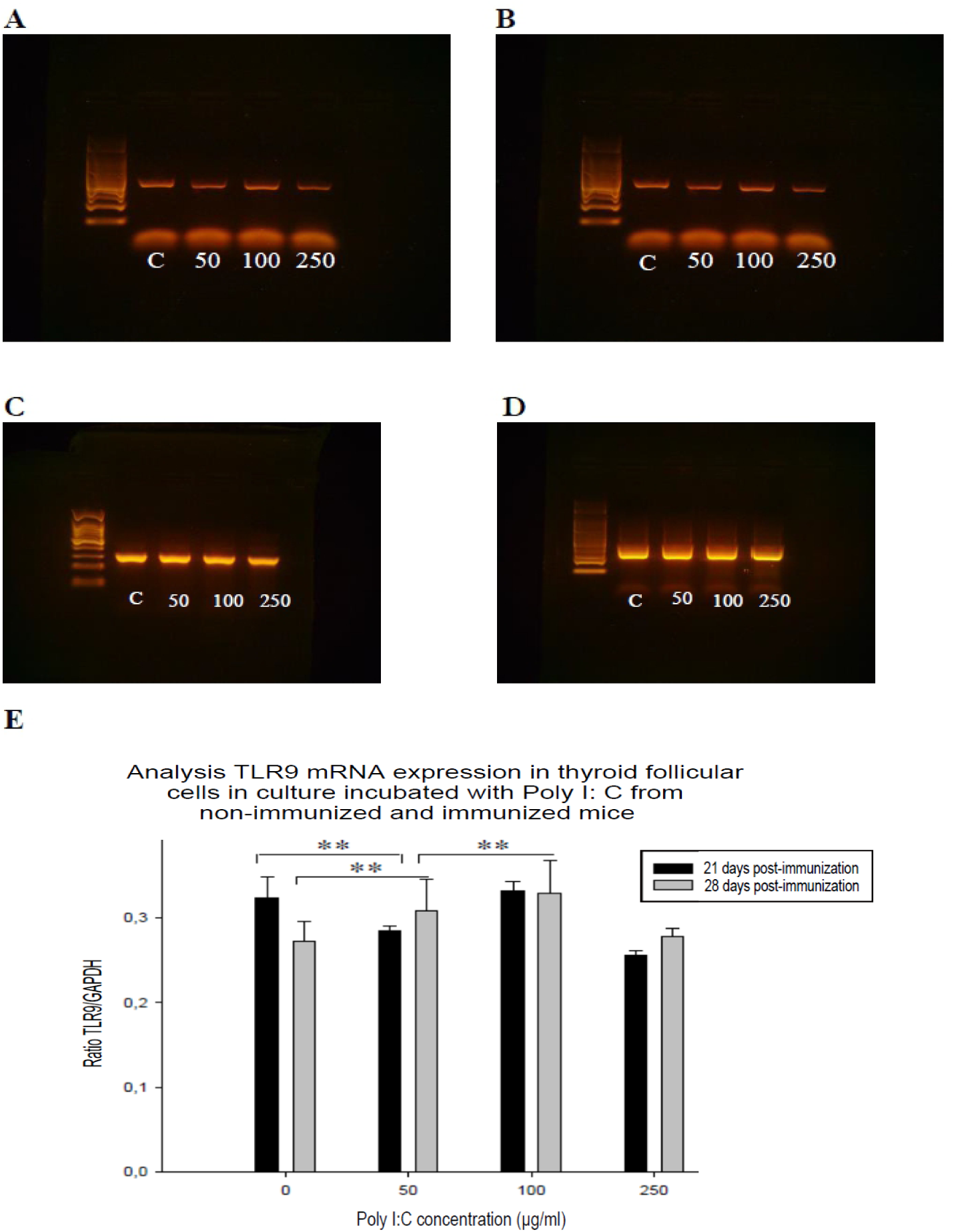
Agarose gels showing PCR-amplified fragments from TLR9 mRNA (417 bp) (**A** and **B**) and GAPDH (274 bp) (**C** and **D**) in thyroid follicles from a pool of 3 mice developed for each of the conditions: non-immunized (**A** and **C**) and immunized (**B** and **D**); which were incubated in cell culture with Poly I: C for 8 hours. **E:** Plot showing TLR9 mRNA expression ratio GAPDH-normalized. C = Control, 50 = Poli I:C 50 ug / ml, 100 = Poli I:C 100 mg/ml, 250 = Poli I:C 250 mg/ml. n = 3. p <0.05 (*), p> 0.05 (**).

Figure 5 refers to the cytokine IL-1. It is observed for both experimental groups an increased mRNA expression relative to control in thyroid follicles when is exposed to either Poly I:C concentration. It can be seen that the basal expression level (control) is markedly lower in the non-immunized animals compared to those immunized and the stimulatory effect is more pronounced in the former. Moreover, the immunized animals for all concentrations show a higher mRNA expression level respect to those non-immunized.

**Figure 5:**
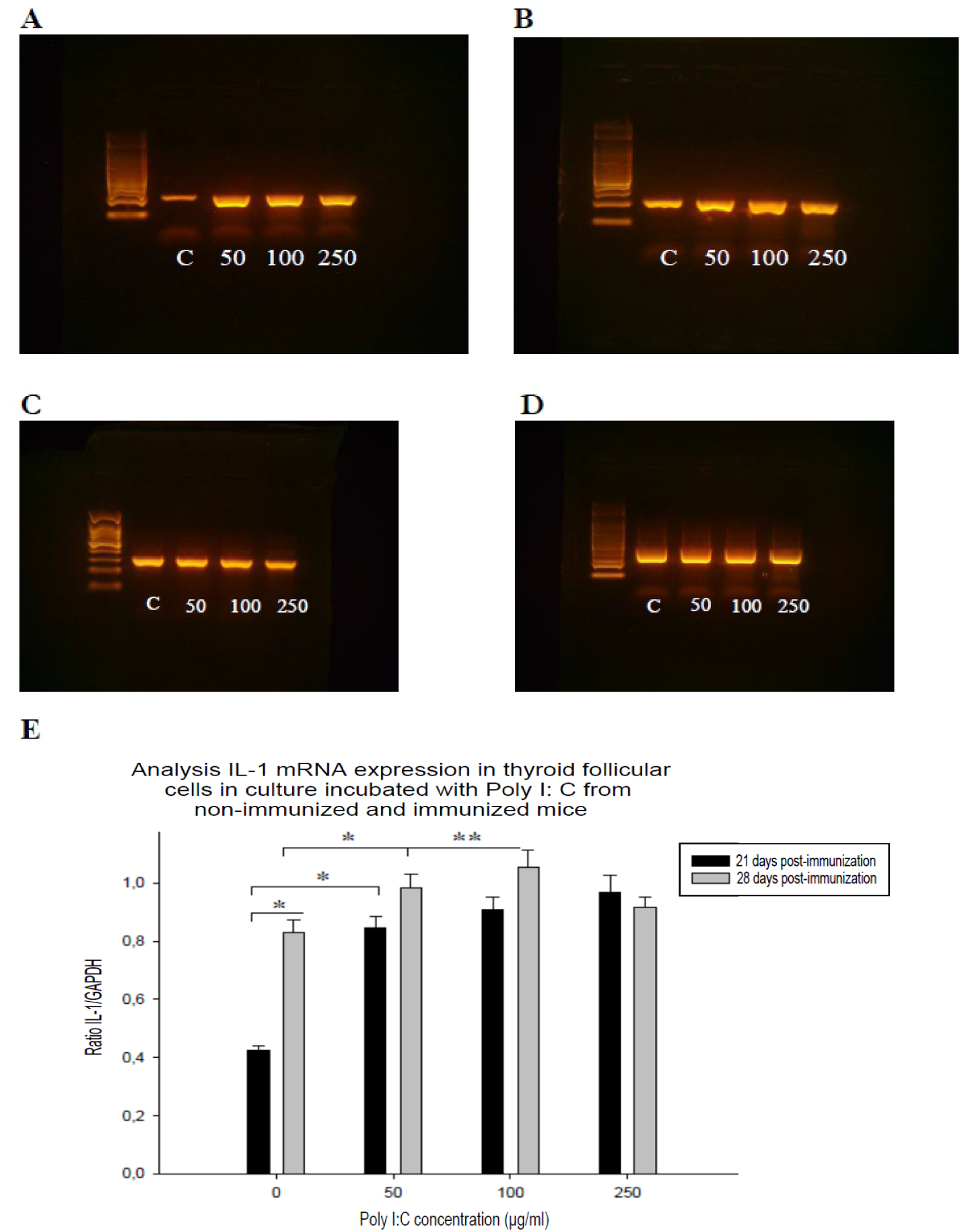
Agarose gels showing PCR-amplified fragments from IL-1 mRNA (200 bp) (**A** and **B**) and GAPDH (274 bp) (**C** and **D**) in thyroid follicles from a pool of 3 mice developed for each of the conditions: non-immunized (**A** and **C**) and immunized (**B** and **D**); which were incubated in cell culture with Poly I: C for 8 hours. **E:** Plot showing IL-1 mRNA expression ratio GAPDH-normalized. C = Control, 50 = Poli I:C 50 ug / ml, 100 = Poli I:C 100 mg/ml, 250 = Poli I:C 250 mg/ml. n = 3. p <0.05 (*), p> 0.05 (**).

In Figure 6 the effect on INF-beta mRNA is evaluated, where exposure of thyroid follicles from immunized animals at the lowest Poly I: C concentration (50 ug/ml) animals is sufficient to achieve a marked increase in the expression levels of this cytokine compared to the control, reaching its greatest intensity at higher concentrations. This finding is observed less markedly in non-immunized animals, where Poly I: C has a minor effect. Again, here we see that the level of basal expression (control) and for each concentration is lower in non-immunized animals compared to those immunized.

**Figure 6:**
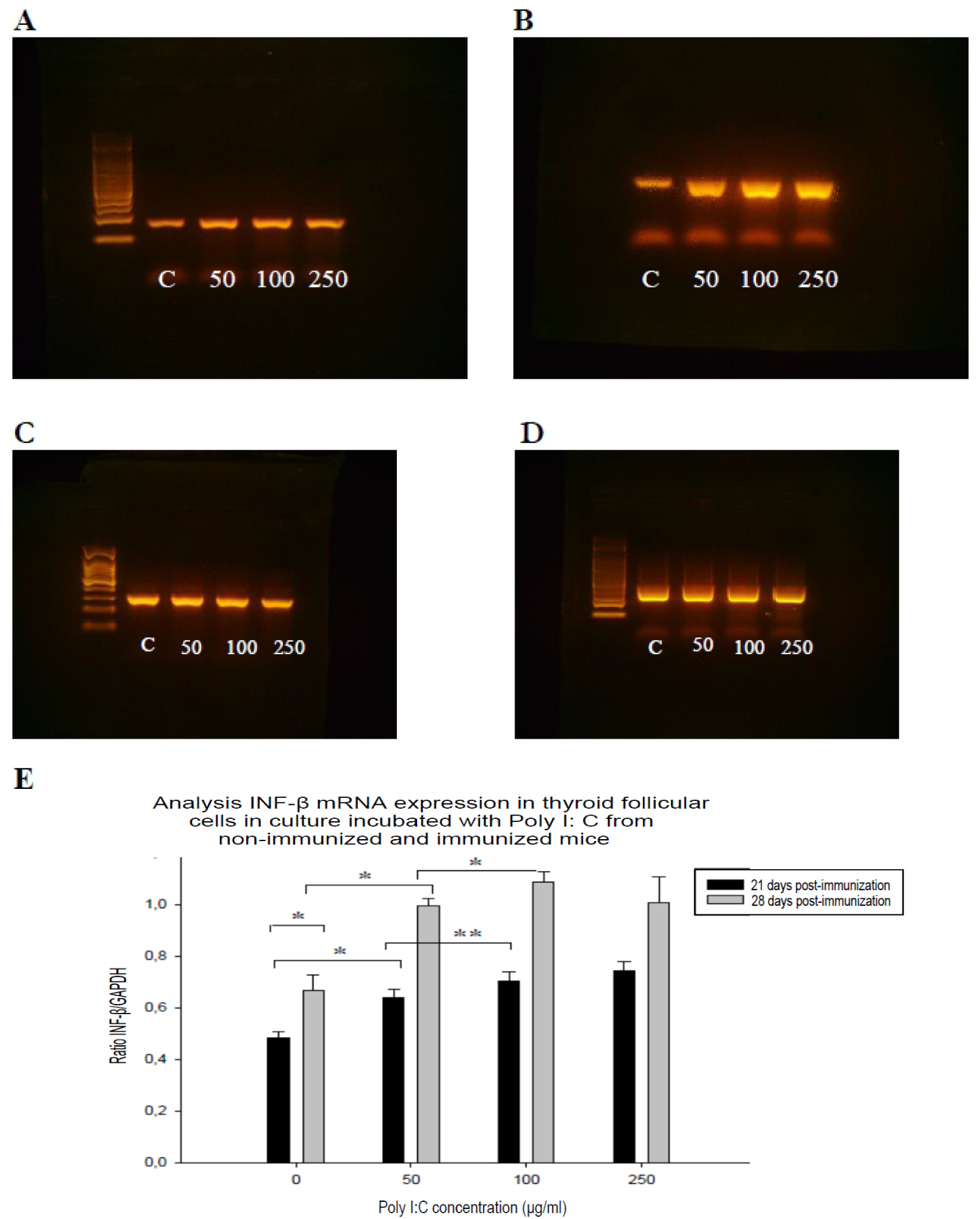
Agarose gels showing PCR-amplified fragments from INF-β mRNA (163 bp) (**A** and **B**) and GAPDH (274 bp) (**C** and **D**) in thyroid follicles from a pool of 3 mice developed for each of the conditions: non-immunized (**A** and **C**) and immunized (**B** and **D**); which were incubated in cell culture with Poly I: C for 8 hours. **E:** Plot showing IL-1 mRNA expression ratio GAPDH-normalized. C = Control, 50 = Poli I:C 50 ug / ml, 100 = Poli I:C 100 mg/ml, 250 = Poli I:C 250 mg/ml. n = 3. p <0.05 (*), p> 0.05 (**).

Figure 7 it is shown differences in the TNF mRNA level expression depending on the synthetic analogue concentrations used. For both experimental groups, Poly I: C 50 and 100 ug/ml on thyroid follicles induces an increase in the mRNA level expression; however, for 250 ug/ml it is produced a marked decrease, it is so pronounced in non-immunized mice that reaches much lower values respect to control. On the other hand, the immunized animals for all concentrations show a higher expression level compared with the non-immunized mice.

**Figure 7:**
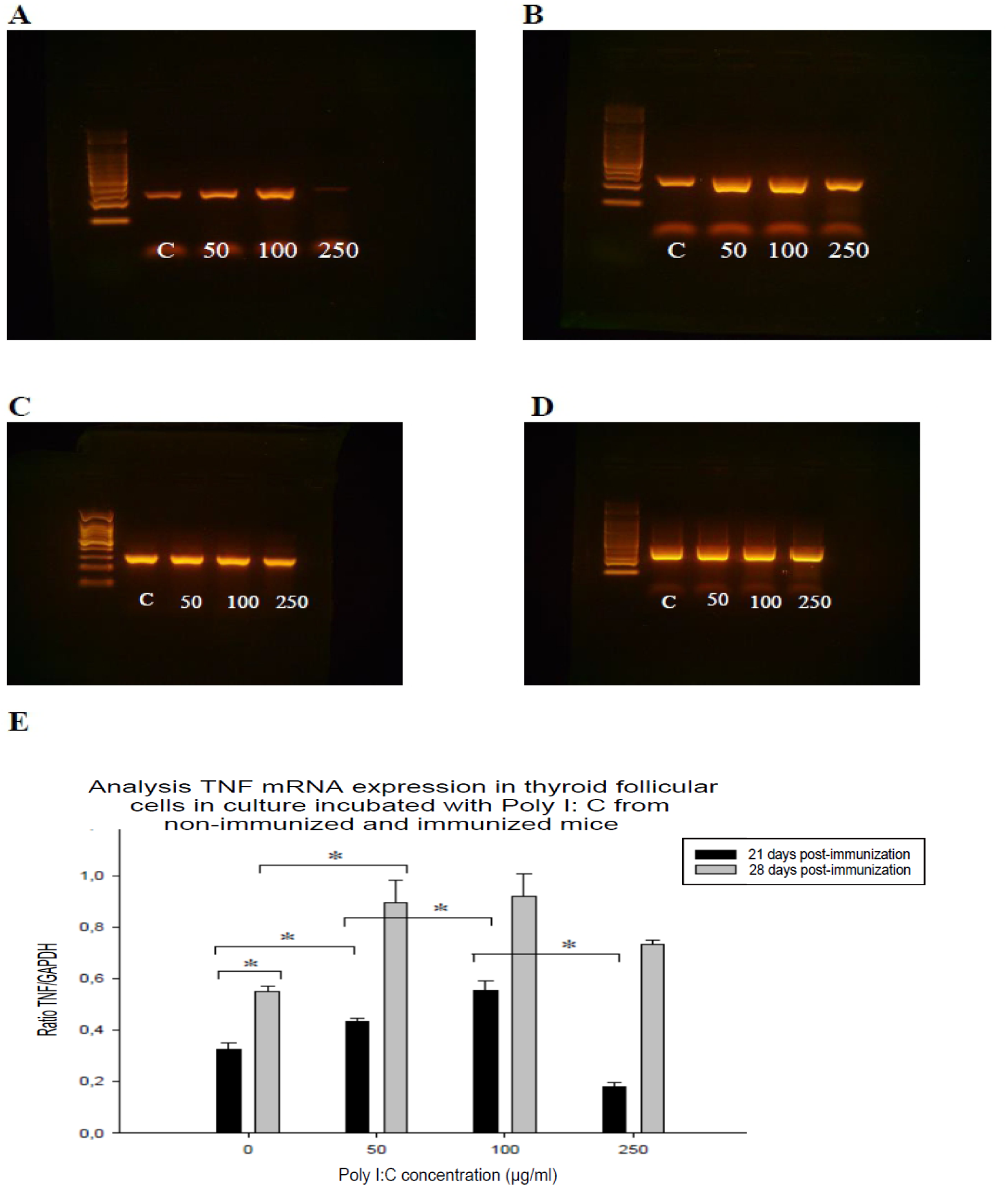
Agarose gels showing PCR-amplified fragments from TNF mRNA (235 bp) (**A** and **B**) and GAPDH (274 bp) (**C** and **D**) in thyroid follicles from a pool of 3 mice developed for each of the conditions: non-immunized (**A** and **C**) and immunized (**B** and **D**); which were incubated in cell culture with Poly I: C for 8 hours. **E:** Plot showing IL-1 mRNA expression ratio GAPDH-normalized. C = Control, 50 = Poli I:C 50 ug / ml, 100 = Poli I:C 100 mg/ml, 250 = Poli I:C 250 mg/ml. n = 3. p <0.05 (*), p> 0.05 (**).

## DISCUSSION

Hashimoto’s thyroiditis is the most common tissue specific autoimmune diseases in humans, characterized by lymphocytic infiltration of the thyroid and production of a high amount of auto-anti-peroxidase, thyroglobulin and anti-TSH-R which generate destruction progressive organ. To study, an experimental disease model called experimental autoimmune thyroiditis, which is induced by parenteral administration in mice murine thyroglobulin is used.

A high proportion of autoantibodies commonly associated with autoimmune diseases (systemic lupus erythematosus, scleroderma, Sjögren’s syndrome, uveitis, etc.), which react with the DNA, RNA or macromolecular complexes containing DNA or RNA is observed. (Marshak-Rothstein, 2006), a process that would take place by activating TLRs with subsequent type I interferons and proinflammatory cytokines expression and adaptive immune response induction. Therefore, assessing overexpression of certain TLR receptors as TLR3 and TLR9 is relevant, whereas their ligands are present in infectious agents, a very important factor as there is a link between certain infections and autoimmunity. In the 90s it was suggested that viral activation of autoimmunity may result from a local tissue infection, an abnormal increase in the expression of MHC genes, autoantigen presentation to immune cells and T cell activation (Harii et al., 2005). An example of an infectious agent that may be involved in autoimmunity induction is the association Foamy Virus infection with Quervain’s thyroiditis.

*In vitro* studies showed that TLR3 agonist, Poly I:C, increased the severity of experimental autoimmune uveitis in association with increased Th1/Th17 levels in mice, delayed type hypersensitivity responses and Ag-specific T cell proliferation (Xiangrong R et al., 2011). With regard to TLR9, studies in animals deficient in this receptor have been controversial and it is unclear whether it plays a key role in the development of autoimmune responses.

In this article, experimental autoimmune thyroiditis was induced in Rockefeller female mice because they respond well to immunization with thyroglobulin. The antithyroglobulin antibody titers were determined after treatment with Poly I:C to evaluate possible differences in the responses to a viral counterpart among non-immunized and immunized mice.

This study planned to analyze the mRNA TLR3 and TLR9 expression levels in thyroid follicular cells from mice following *in vitro* stimulation with Poli I:C and determining its ability to enhance cytokine expression capable to intervene in the development and progression of the disease, such as INF-β, TNF and IL-1. In addition, we evaluated whether Poly I: C has an adjuvant effect after stimulate TLR3 involved in the experimental autoimmune thyroiditis induction, through increased production of anti-thyroglobulin antibodies.

For mRNA TLR3 and TLR9 expression levels and its subsequent mRNA proinflammatory cytokines expression (IL-1, TNF and IFN-beta), it expected to be of different magnitude in the immunized animals, as it has been seen that they show mRNA expression levels higher that animals non-immunized, as was seen with TLR3 in Hashimoto’s thyroiditis in humans (Harii et al., 2005).

Poly I: C is the synthetic dsRNA analog and is an immunostimulant to be TLR3 ligand and in these conditions simulate a thyroid follicles viral infection. The expression of said mRNA is performed in thyroid follicular cells from non-immunized and immunized animals. Whereas EAT induction causes morphological changes in the thyroid gland and immune cells migration Whereas APR induction causes morphological changes in the thyroid gland and migration of immune cells that may affect the test results, we isolate thyroid follicles in primary culture and these were stimulated with microbial analog. This proved not induce morphological changes in the follicles during incubation for 18 hours, allowing more reliably study the effect of these analogues on mRNA TLR3, TLR9, IL-1, INF-beta and TNF expression level in thyroid follicles from immunized and non-immunized animals.

Previous studies in our laboratory showed detectable amounts of TLRs (mRNA and protein) in thyrocytes from control animals and increased expression over time in this receptor in mice with EAT. Furthermore, such experiments showed that no significant differences in mRNA TLR3, TLR9, IL-1, INF-beta and TNF expression when incubated with analogues for different times. Therefore, we opted for a period of 8 hours.

Evaluating the mRNA expression in thyroid follicular cells from non-immunized and immunized mice it can be seen in most conditions an increase in expression levels after stimulation with Poly I:C. It was noticeable that this increase is more intensely developed in immunized as compared with non-immunized mice even observed differences from baseline in both groups. This suggests that animals with EAT have a TLR3, TLR9 and inflammatory cytokines basal expression level higher, may contribute to cause or maintain a chronic damage of the gland when the autoimmune disease is already established. Moreover, this might cause same thyroid follicular cells from diseased individuals are more sensitive to infectious stimulus and be able to quickly activate signaling cascades via TLRs so leading to express these receptors and consequently increased proinflammatory cytokines, may with thereby increasing the severity of symptoms and perpetuate it in time.

On the other hand, it could also be noted that curiously certain high Poly I: C concentrations generated the opposite effect: a reduction in expression levels, a phenomenon that occurred with 250 ug/ml on mRNA TLR3 and TNF and may indicate that activation of the TLR signaling pathway occurs within a range of ligand concentration, thus higher values not induce or inhibit the effect. Also shows that low concentrations were able to induce a significant impact on several conditions, in some cases being equal to or greater magnitude than those resulting from higher concentrations, can be observed with Poly I: C on mRNA TLR3 and TNF.

With the above, we note that in experimental autoimmune thyroiditis, infectious agents may have an adjuvant effect by stimulating TLR receptors in thyroid follicular cell inducing increased in TLR and proinflammatory cytokines expression levels, may be able to intervene in the development and progression of autoimmune thyroiditis.

